# Scalable Guide Tree Construction Using Quantum Annealing for Multiple Sequence Alignment

**DOI:** 10.1101/2024.11.30.626202

**Authors:** Youngjun Park, Juhyeon Kim, Joonsuk Huh

**Author notes:** These authors contributed equally to this work (Youngjun Park, Juhyeon Kim).

## Abstract

Multiple sequence alignment (MSA) reveals homology in biological sequences, which is crucial for phylogenetics, medicine, and molecular biology. Many heuristic MSA algorithms use guide trees to determine sequence alignment order, but finding an optimal guide tree is an NP-hard problem. Conventional guide tree algorithms are greedy heuristics designed for better scalability at the cost of accuracy. By utilizing quantum algorithms, such as quantum annealing, we can overcome the problem of local minima that occurs in greedy methods. We propose a scalable guide tree algorithm that achieves scalability through quantum annealing. The theoretical foundations of our method are both minimum evolution and molecular clock. Unlike classical greedy approaches, we directly map the minimum evolution problem to a traveling salesperson problem (TSP), which quantum annealing can solve efficiently. The TSP solution enables the guide tree to be constructed in linear time using the molecular clock. Even with only a single sample from a D-Wave hybrid solver, our guide tree generally performed comparably to classical trees on the BAliBASE 3.0 benchmark. With a single iterative refinement, no statistically significant performance differences were found between our method and classical guide trees. Our scalable guide tree algorithm is practical in the sense that only a single sampling can be enough for constructing a good guide tree. Further performance analysis requires larger-scale benchmark tests. Fortunately, rapid advances in quantum hardware may soon enable these tests. While the practical application of quantum algorithms in bioinformatics has been relatively overlooked, this study highlights the potential of quantum algorithms for targeting computational bottlenecks in the field.

## I. INTRODUCTION

Multiple sequence alignment (MSA) is a fundamental method for interpreting genomic and proteomic data from various organisms. It plays a pivotal role in identifying *homologies* among biological sequences [1, 2]. Based on homology, phylogenetic trees can be constructed for various research areas, including disease treatment [3], pathogenesis [4], drug target identification [5], protein-protein interactions [6], enzyme research (e.g., Cas9 protein [7]), and the prediction of protein structure [8–10] and function [5].

Unfortunately, computing an optimal MSA is an NP-hard [11, 12] combinatorial optimization problem which becomes intractable even at a moderate problem size. Therefore, MSA tools employ combinations of heuristic algorithms, including progressive alignment, iterative refinement, consistency-based schemes, profile hidden Markov models, and phylogeny-aware methods [1, 13– 26]. Most of the MSA tools [1, 13–17, 19–26] perform progressive alignment that requires a *guide tree*.

For progressive alignment, *N* (*N* − 1)*/*2 pairwise alignments using the Needleman-Wunsch [27] or Smith-Waterman algorithm [28] are performed to compute rel-ative distances between pairs of sequences [29], where *N* is the number of sequences. A guide tree is constructed solely from the distances to determine the sequence alignment order [30]. Guide tree construction algorithms are distance-based algorithms which should be distinguished from other phylogenetic inference methods, including maximum-likelihood [31], parsimony [32], and Bayesian inference [33]. While the guide tree algorithms require only a distance matrix as input, the phylogenetic inference methods are based on specific mathematical models independent of the distance matrix.

Guide trees significantly influence the quality of MSA results [34]. However, finding an optimal tree is an NP-hard problem [35– Many guide tree algorithms are hierarchical agglomerative clustering (HAC) [39, 40], which is a bottom-up heuristic with low computational complexity. Neighbor-joining (NJ) [41] and unweighted pair group method with arithmetic mean (UPGMA) [42] are the most widely used guide tree algorithms. They first compute a full distance matrix, where parallel computing can effectively reduce the runtime as each distance can be independently calculated. Extensive research [16, 22, 43– 46] has been conducted to improve the time complexity of the NJ and UPGMA. The best implementations [22, 46] achieve a time complexity of *O*(*N* ^2^) for a given distance matrix; however, the complexity is still considered unscalable. PartTree [47] and a modified version [1] of mBed [48] are scalable methods with *O*(*N* log *N*) time complexity. However, these methods calculate a portion of a distance matrix and may yield less accurate trees. Moreover, PartTree is generally incompatible with other MSA tools [48].

Recently, there has been growing interest in utilizing quantum algorithms for solving NP-hard problems in bioinformatics [49–54]. Many NP-hard combinatorial optimization problems can be formulated as an Ising Hamiltonian, which encodes the solution as its ground state. Quantum annealing [55] and the quantum approximate optimization algorithm (QAOA) [56] are quantum algorithms specialized for efficiently solving combinatorial optimization problems expressed as the Ising Hamiltonian. Quantum annealing is based on the quantum adiabatic theorem, which states that if a quantum system undergoes a slow evolution, an eigenstate of the initial system evolves into the corresponding eigenstate of the final system [57]. We initialize the system in the ground state of a transverse-field Hamiltonian, which is a uniform superposition of all possible combinatorial instances. The superposition is easy to prepare, and we can slowly evolve the initial Hamiltonian into the target Ising Hamiltonian. During the evolution, quantum tunnelling [58] allows the system to approach the global minimum by penetrating potential barriers. Unwanted excitation from the ground state can be mitigated by choosing a gapped path [59] or introducing counterdiabatic driving to the evolution [60, 61]. By measuring the system qubits after the evolution, we can obtain an approximate solution to the problem. While quantum annealing evolves a system in continuous time, the QAOA discretizes the time into steps due to its implementation on digital quantum computers [62]. It is worth noting that quantum annealing has scaling advantages over simulated annealing [63] and path-integral Monte Carlo [64].

In this research, we present a scalable guide tree algorithm using quantum annealing, which shows generally comparable performance to classical guide tree algorithms in BAliBASE 3.0 benchmark test [65] even with a single sampling for quantum annealing. This algorithm is compatible with any MSA tool that accepts distance-based guide trees. Our approach is based on two theoretical principles in guide tree construction: minimum evolution (ME) [66–69] and molecular clock (MC) [70– 73]. The ME assumes that an optimal tree has a minimum sum of branch lengths. In contrast to conventional greedy approaches to the ME problem [68], we directly solve it using quantum annealing by mapping the ME problem to the traveling salesperson problem (TSP) [74]. The QAOA can also solve the TSP, but in this study, we primarily focus on the scaling advantage of quantum annealing [63, 64]. The TSP solution is a circular order of sequences that can be used for building a tree with a minimum sum of branch lengths. Using the circular order, our algorithm applies the MC to construct a guide tree with positive branch lengths. This step in our algorithm has a linear time complexity of *O*(*N*), which is quadratically faster than the *O*(*N*^2^) complexity of conventional guide tree algorithms.

Recent studies have reported quantum annealing and the QAOA approaches to MSA [51–53]. Nevertheless, these algorithms are limited to small-sized toy problems and cannot insert gaps in the longest sequence. Previous research on MSA and tree construction based on the TSP [74–76] has been conducted. However, these studies did not specify how to determine and ensure positive branch lengths. They even omitted assigning branch lengths on guide trees for the MSA process. All of these issues are addressed in our proposed algorithm.

The rest of this paper is organized as follows. Section II introduces our guide tree construction method. First, in Subsection II A, we define minimum evolution with molecular clock (MEMC) principle and explain how it addresses issues in previous research. Then, in Subsection II B, we introduce our scalable guide tree algorithm, referred to as the MEMC algorithm, based on this principle. A parallel computing scheme for the distance matrix computation is detailed in Appendix A. In Section III, we present the BAliBASE 3.0 benchmark test results, comparing the MEMC algorithm with classical guide tree methods. Data preparation details and supplementary data are provided in Appendix B and Appendix C, respectively. Finally, we discuss our findings in Section IV and conclude the paper in Section V.

## II. METHODS

### A. The MEMC Principle

We establish the MEMC for constructing guide trees, which is stated as follows:

#### **Principle 1**.

*(MEMC principle) In the optimal guide tree for a given set of sequences, (1) the sum of branch lengths is minimized, and (2) the branch lengths from the root to each leaf node are equal*.

The first part of this principle is based on the ME, which is then followed by the application of the MC with a modification from this work.

#### 1. The ME in the MEMC Principle

The ME considers a tree to be optimal if it represents a minimum number of evolutionary events [41, 66, 67]. Because each branch length is proportional to the number of evolutionary events, the optimal tree has the minimum sum of branch lengths. However, finding the optimal tree according to the ME is an NP-hard optimization problem [37, 38]. The NJ takes a greedy approach to the ME problem, which results in *O*(*N*^2^) time complexity in its best implementation [46].

In this work, we solve the ME problem using a non-greedy approach known as the circular sum (CS) method [74, 75]. Before describing the method in detail, we introduce the necessary notation. A circular order *C* of input sequences is an order in which each sequence *s*_*i*_ appears exactly once, for example, *C* = [*s*_3_, *s*_1_, *s*_4_, *s*_2_]. The sequence at the *n*th position of *C* is denoted by *C*_*n*_, such that for this example, *C*_1_ = *s*_3_, *C*_2_ = *s*_1_, *C*_3_ = *s*_4_, and *C*_4_ = *s*_2_. *d*(*s*_*i*_, *s*_*j*_) denotes the distance between sequences *s*_*i*_ and *s*_*j*_ in a distance matrix.

The CS is the sum of distances between each pair of sequences that are consecutive in a given circular order 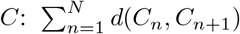. As shown in Fig. 1, this sum is independent of tree topologies and is always twice the sum of branch lengths. The ME problem can be alternatively solved by finding the circular order that minimizes half of the CS:

**FIG. 1.**
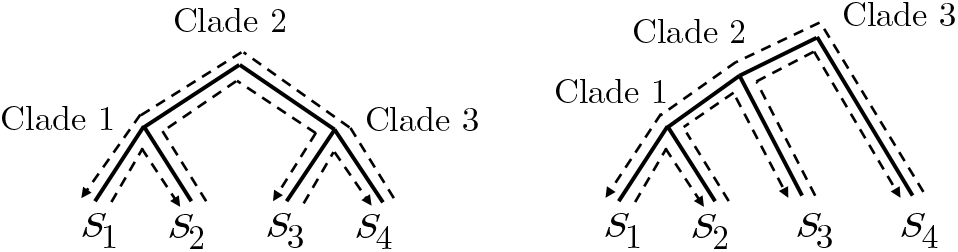
The CS method can calculate the sum of branch lengths independent of tree topologies.

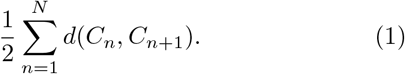

Therefore, the CS method maps the ME problem to TSP, which can be efficiently solved by quantum annealing.

#### 2. The MC in the MEMC Principle

After solving the TSP, we obtain a circular order *C*. We apply the MC to the circular order for determining a tree topology while ensuring positive branch lengths and the scalability of our method. Furthermore, our approach using the MC can avoid several issues with the CS method in the original work [74].

The first issue in the CS is that optimal pairwise alignment scores were used to compute the CS and assign branch lengths to a tree. However, the optimal pairwise alignment scores can be negative for sequences with low similarity. The original work introduces a criterion for determining tree topologies (see 74, Definition 12]), but does not clarify how to assign branch lengths. The criterion is vulnerable to introducing negative branch lengths because a longer distance *d*_*i*_ can be selected earlier than a shorter distance *d*_*j*_ in the algorithm, while there is a branch length computed as *d*_*j*_ − *d*_*i*_ < 0. By repeating *O*(*N*) steps each with *O*(*N*) for the criteria, *O*(*N*^2^) computational time is required, which is not scalable.

In the present work, we use positive distances computed by an MSA tool, where we intend to input our guide trees. The other issues can be avoided by applying the MC. The MC assumes that the rate of molecular evolution is approximately constant in different species [72, 73]. This implies that the distances from the root to each leaf node are the same. If we arrange the leaf nodes according to the circular order *C*, we can apply the MC. We modify the MC not to update a distance matrix, which enables *O*(*N*)-time implementation of our algorithm and ensures assigning positive branch lengths.

### B. MEMC Guide Tree Construction Algorithm

Now we explain the details of our guide tree construction algorithm: the MEMC algorithm, which is based on the MEMC principle (Principle 1). In the first phase of the algorithm, a TSP related to the ME problem is solved by quantum annealing, where we obtain a circular order *C*. We then determine a tree topology and positive branch lengths from *C*, with a linear time complexity, *O*(*N*). This linear time complexity enables scalable guide tree construction. The branch lengths are determined in a similar way to UPGMA algorithm [42]. Figure 2 shows the overall process with a simple example.

**FIG. 2.**
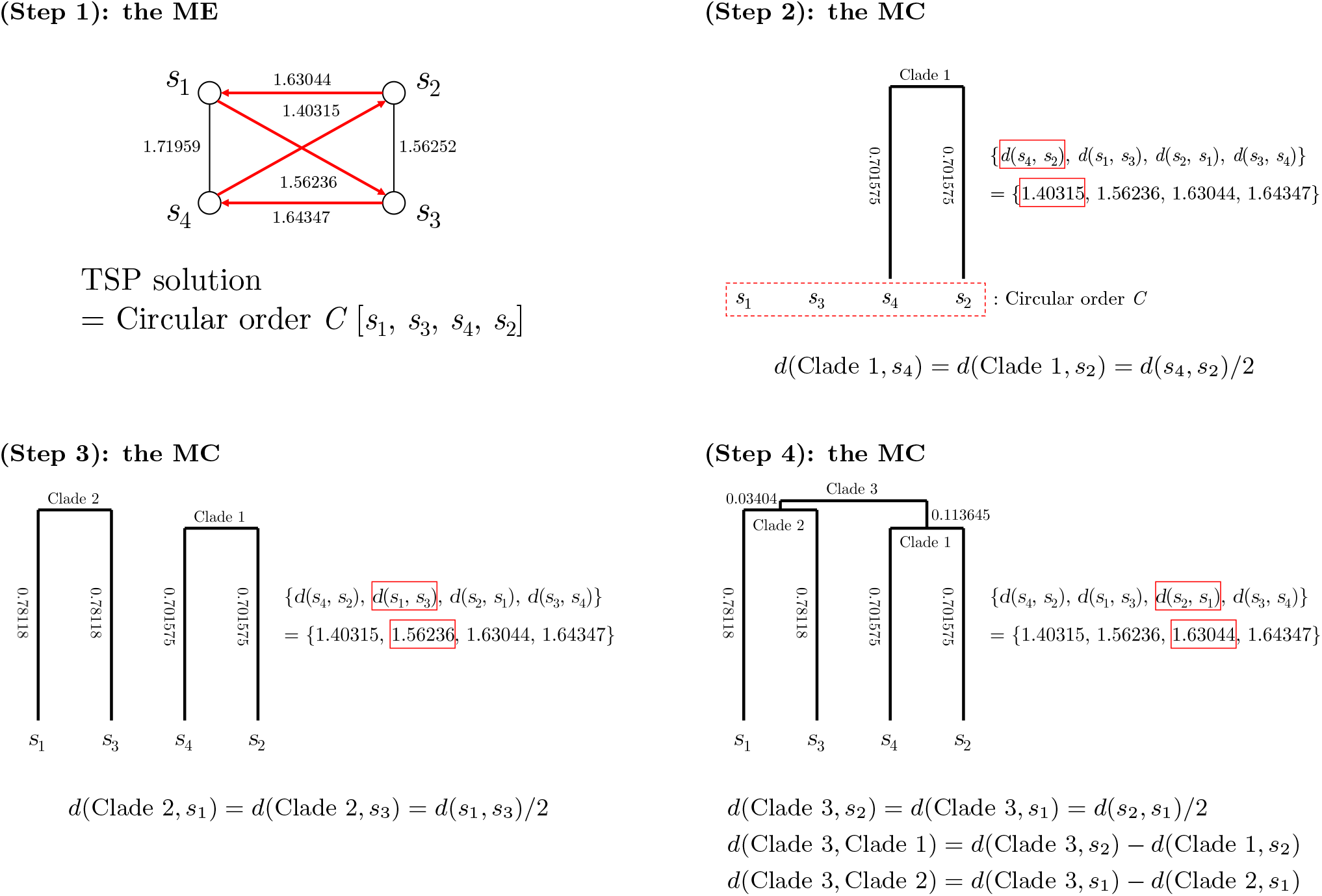
The MEMC guide tree construction algorithm. Step 1 is the first phase, and Steps 2–4 are the second phase of the algorithm. Step 1 finds a circular order *C* from a TSP related to the ME. Then, the distances between consecutive sequences in *C* are saved and sorted in increasing order. The tree topology and positive branch lengths are determined through Steps 2–4 while not breaking the circular order *C*. At each step, the two closest leaf nodes or clades are connected, and the newly generated branch lengths are determined. This step is repeated *N* − 1 times for *N* input sequences until a rooted tree is constructed. Note that the distance matrix is not updated after each clustering.

#### 1. Generating a TSP Graph for Finding a Circular Order

A distance matrix, which can be either given or computed through parallel computing (Appendix A), is first mapped to a TSP graph to obtain a circular order *C*. Each node in the graph corresponds to an input sequence *s*_*i*_. The weight of the edge between two nodes representing *s*_*i*_ and *s*_*j*_ is given by the distance *d*(*s*_*i*_, *s*_*j*_). The graph is fully connected, undirected, and satisfies *d*(*s*_*i*_, *s*_*j*_) = *d*(*s*_*j*_, *s*_*i*_) for all *i ≠ j*. An example is shown in Step 1 of Fig. 2. The distances must be calculated using the same metric as the MSA tool that will receive the guide tree. Using a different distance measure can lead to inaccurate results by misinterpreting the evolutionary closeness between sequences.

#### 2. Finding the Circular Order Using a Quantum Annealer

A quantum annealer is used to find the solution to the TSP from the graph. To formulate the TSP as a binary model, we define a binary variable *b*_*i,p*_ ∈ {0, 1} . A binary variable *b*_*i,p*_ is defined to take the value of 1 if the node representing *s*_*i*_ is selected at the *p*-th step of a circular order *C*, and 0 otherwise:

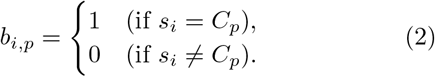

Using *b*_*i,p*_, the TSP can be formulated as the following Ising model [77]:

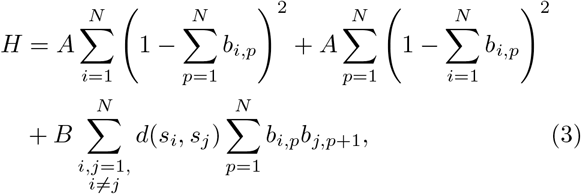

where *A* is a positive constant, and 0 *< B* · max_1≤*i,j*≤*N*_ (*d*(*s*_*i*_, *s*_*j*_)) *< A*. The node selected at the (*N* + 1)th step is the same as the initial node. The condition *B* · max_1≤*i,j*≤*N*_ (*d*(*s*_*i*_, *s*_*j*_)) *< A* strictly excludes non-Hamiltonian cycles from solutions. Using the binary-to-spin mapping *b* ↦ (1 + *σ*)*/*2, where *σ* ∈{−1, 1}, we obtain the Ising Hamiltonian for the TSP from the Ising model. Quantum annealers can obtain the approximate ground state of the Ising Hamiltonian with better scaling in time-to-solution than simulated annealing [63] and path-integral Monte Carlo [64]. Therefore, using quan-tum annealing in the MEMC algorithm enables scalable guide tree construction. Additionally, the QAOA can be utilized to find the approximate ground state of the Ising Hamiltonian, which enables universal quantum computation for the MEMC algorithm [56, 78]. The binary representation of the ground state corresponds to the solution of the TSP. For instance, if *b*_1,1_, *b*_3,2_, *b*_4,3_ and *b*_2,4_ are ones and the others are zeros, then *C* = [*s*_1_, *s*_3_, *s*_4_, *s*_2_] (Step 1 in Fig. 2).

#### 3. Determining the Tree Topology and Branch Lengths

We determine the topology and branch lengths of the guide tree from *C* using the MC. First, each leaf node is assigned to represent an input sequence in the order of *C*. Second, the *N* distances between consecutive pairs of sequences in *C* are saved as a set 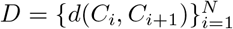. The set *D* is sorted in an increasing order of distance using radix sort, which requires a worst-case time complexity of *O*(*kN*) [79], where *k* is the number of significant digits in the distances. The number of significant digits *k* is a constant, which is at most six to seven digits. Therefore, *k* ≪ *N* in practical cases. Finally, using the sorted set *D*, the corresponding nodes or clades are identified and connected, and their branch lengths are computed. This part of the algorithm is illustrated in Steps 2–4 of Fig. 2. Let *C* = [*s*_1_, *s*_3_, *s*_4_, *s*_2_] be the solution of the TSP. From *C*, we have the set of distances *D* = ≪*d*(*s*_1_, *s*_3_), *d*(*s*_3_, *s*_4_), *d*(*s*_4_, *s*_2_), *d*(*s*_2_, *s*_1_) }. A sorted set *D*^*′*^, for example, is *D*^*′*^ = {*d*(*s*_4_, *s*_2_), *d*(*s*_1_, *s*_3_), *d*(*s*_2_, *s*_1_), *d*(*s*_3_, *s*_4_)}.

The first element *d*(*s*_4_, *s*_2_) of the sorted set *D*^*′*^ is the minium distance in the set *D*. Two leaf nodes representing *s*_4_ and *s*_2_ are therefore connected to a new clade (Clade 1) as shown in Step 2 of Fig. 2. At this step, we apply the MC to make the new clade equidistant from the connected leaf nodes, such that the branch lengths are set as *d*(Clade 1, *s*_4_) = *d*(Clade 1, *s*_2_) = *d*(*s*_4_, *s*_2_)*/*2.

After connecting either nodes or clades, classical HAC algorithms update distances to reflect new relationships between the new clade and other existing nodes and clades. If the distances are updated in the MEMC algorithm, a new TSP will have to be solved to find a circular order for the updated distance matrix. This requires solving a total of *O*(*N*) different TSPs. Therefore, in our algorithm, distances are not updated because the circular order obtained using quantum annealing can be considered as a good approximation to the global minimum of the sum of branch lengths.

The process is repeated without updating the distances in the sorted set *D* until all leaf nodes are connected to a single tree (Steps 3 and 4 in Fig. 2). Note how branch lengths are computed when two clades are connected to a new clade (Step 4 in Fig. 2). The branch length between the new clade and one of the connected clades is the difference between the branch lengths of these clades. Thus, *d*(Clade 3, Clade 1) = *d*(Clade 3, *s*_2_) − *d*(Clade 1, *s*_2_), and *d*(Clade 3, Clade 2) = *d*(Clade 3, *s*_1_) − *d*(Clade 2, *s*_1_), where *d*(Clade 3, *s*_2_) = *d*(Clade 3, *s*_1_) = *d*(*s*_2_, *s*_1_)*/*2. This computation guarantees assigning positive branch lengths because *d*(*s*_2_, *s*_1_) *> d*(*s*_1_, *s*_3_) *> d*(*s*_4_, *s*_2_) is ensured by the sorted distance set *D*. A similar computation is applied when a clade and a leaf node are connected to a new clade.

To ensure efficient implementation of the algorithm, a disjoint-set data structure combined with two optimization techniques, namely union by rank and path compression, is employed [79]. The process begins by creating a disjoint-set data structure for *N* leaf nodes, which has a time complexity of *O*(*N*). Path compression accelerates the identification of the clade to which a leaf node belongs, and union by rank efficiently joins two subtrees. Both operations have a time complexity of *O*(*α*(*N*)), where *α*(*N*) is a very slowly growing function known as the inverse Ackermann function. Our algorithm executes a constant *O*(1) number of these operations at each step, leading to a total time complexity of *O*(*Nα*(*N*)) over *O*(*N*) steps. Note that *α*(*N*) ≤ 4 for any practical value of *N* [79]. In conclusion, the worst-case time complexity of the proposed algorithm, including the sorting step, can be considered linear: *O*(*N* (*k* + *α*(*N*))) ≈ *O*(*N*).

## III. RESULTS

In this section, we present BAliBASE 3.0 [65] benchmark test results to compare the performance of MEMC guide trees with conventional guide trees. We input different guide trees to MAFFT L-INS-i (ver. 7.526) for aligning each BAliBASE test case, and compute the sum of pairs (SP) and total column (TC) scores of the MSA results. However, the MEMC algorithm is also compatible with other MSA tools and is not limited to MAFFT. Our MEMC guide trees achieve performance comparable to conventional trees, even when using a single sample from quantum annealing and one iterative refinement in MAFFT L-INS-i. We used a D-Wave quantum-classical hybrid solver for solving the TSP. The data preparation details for this section are described in Appendix B.

### A. Performance Evaluation

We evaluate the performance of the MEMC guide trees against classical distance-based HAC guide trees, including complete linkage (CL), flexible, modified UPGMA (mUPGMA), NJ, single linkage (SL), UPGMA, UPGMC, Ward, WPGMA, and WPGMC [80]. Mupgma is the default guide tree algorithm in MAFFT [81]. For the flexible [82] guide tree algorithm, we arbitrarily set its parameter *β* = 0.5. We set the minimum branch length to 0.00001 to prevent negative branch lengths in the classical guide trees, while five significant digits are used for branch lengths.

BAliBASE 3.0 is a widely used protein MSA benchmark for evaluating the performance of MSA tools and guide trees [83]. There are 218 test cases, which are categorized into six different groups: BB1-1 (38 families), BB1-2 (44 families), BB2 (41 families), BB3 (30 families), BB4 (49 families), and BB5 (16 families).

For the MEMC algorithm, each TSP was solved using a single shot from the quantum annealer. The single sampling is for simulating a realistic situation where rapid guide tree construction by limiting the number of shots is prioritized. Therefore, the results presented here highlight the promising performance of our method even with this minimal sampling setting.

#### 1. *Results with a Minimum Iterative Refinement:* maxiterate 1

MAFFT L-INS-i is an iterative, progressive, and consistency-based alignment method. It requires at least a few iterative refinements to enhance the accuracy of progressive alignment. We experimented with `maxiterate 1` to minimize the effect of the MSA algorithm on the performance of guide trees.

Fig. 3 summarizes the results of `maxiterate 1`. The complete dataset for Fig. 3 is available in Tables C3 and C4 of Appendix C. As shown in these tables, all of the p-values are greater than 0.05 and F-values are less than the critical F-value 1.83. Therefore, we found no statistically significant difference between the scores of the MEMC guide trees and those of the conventional ones at the *p* = 0.05 level. These results support that our guide tree algorithm can work well in an extreme situation using only a single shot and a single refinement.

**FIG. 3.**
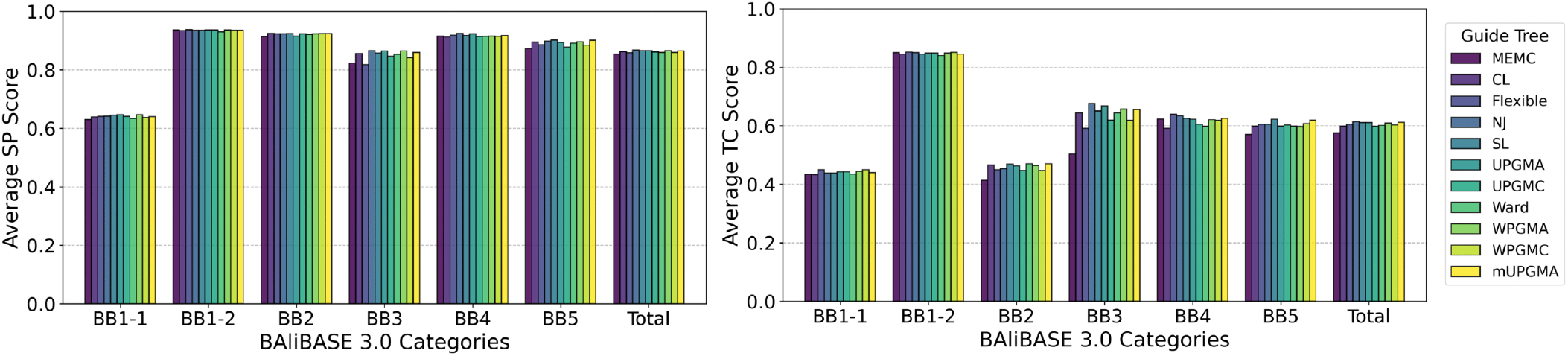
MEMC guide trees show comparable performance to conventional methods on the BAliBASE 3.0 benchmark. The upper and lower charts show the average SP and TC scores, respectively. All alignments were generated using MAFFT L-INS-i (ver. 7.526) with `maxiterate 1`. Each MEMC tree was constructed using a single sample from the D-Wave hybrid solver. There were no statistically significant differences between the methods for either scores (for all cases, *p >* 0.05). Full data are available in Tables C3 and C4 (Appendix C). CL: Complete linkage; mUPGMA: Modified UPGMA; SL: Single linkage.

#### 2. Iterative Refinement Improves Scores for Alignments Using the MEMC Guide Trees

We now turn to analyze how the number of iterations affects the SP and TC scores of the MEMC guide trees. When `maxiterate 0`, MAFFT performs a single progressive alignment using a given guide tree without iterations. `maxiterate 0` thus may directly show the performance of guide trees themselves. However, the results should be carefully interpreted since the results are also affected by the progressive alignment, and iterative refinements are necessary for MAFFT L-INS-i.

Without iterations, the TC scores of the MEMC trees for BB3 (*p <* 0.001) and total (*p* = 0.003) are found to be statistically significantly worse than those of the conventional methods (Tables C1 and C2 of Appendix C). For the other benchmark categories, its performance was comparable to that of the classical methods. We further analyzed these two datasets by Tukey’s HSD test. The TC score of the MEMC algorithm is worse than all the classical methods for the BB3 category (*p <* 0.05). The total TC score of the MEMC trees is significantly worse (*p <* 0.05) than flexible, SL, UPGMC, WPGMC, and mUPGMA, but there are no significant differences (*p >* 0.05) with CL, NJ, UPGMA, Ward, and WPGMA.

Except for the BB3 and total scores, the differences in TC scores between the MEMC and classical guide trees are not statistically significant, which is supported by both *p >* 0.05 and *F <* 1.83. Therefore, we conclude that the low total TC score of our method results solely from the low performance for the BB3 category for the case of `maxiterate 0`.

The BB3 category has the highest average number of sequences per test case (63.17 sequences/file), while the other categories range from 6.87 to 45.59. BB3 also contains two test cases with the two highest numbers of sequences (142 and 140 sequences). Because the size of the tests in BB3 is larger than the others, the hybrid D-Wave solver may find difficulty in obtaining good approximate solutions. As there have been dramatic advances in the performance of the hybrid solver, the results for BB3 will be improved in the near future.

BB3 comprises sequences of similar lengths, which are clustered into several subfamilies [65, 84]. A subfamily is a subset of input sequences that have closer biological relationships (*>* 25% sequence identity) than other sequences. This implies that branch lengths within a subfamily are shorter than those between subfamilies. The MEMC algorithm tends to generate trees with relatively shorter branches adjacent to the root (Fig. 4(a)) than classical algorithms (e.g. UPGMA in Fig. 4(b)). This difference is caused by directly solving the ME problem, which can include long branch lengths to be considered.

**FIG. 4.**
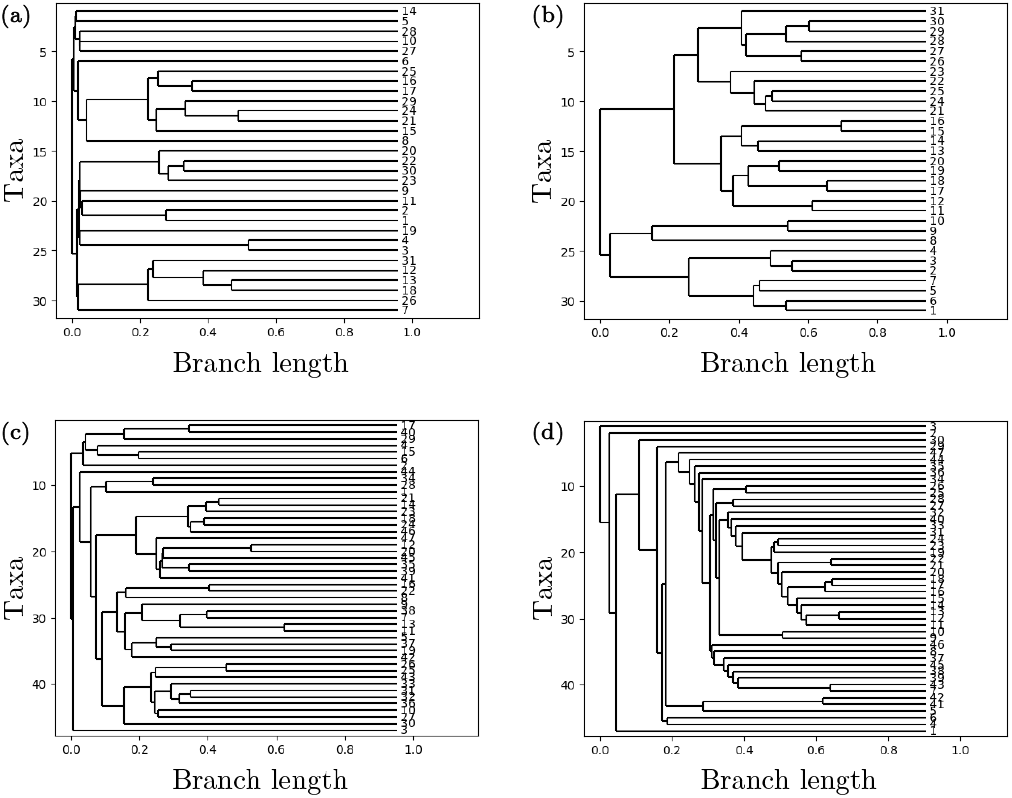
Examples of guide trees. (a) MEMC (BB30002). (b) UPGMA (BB30002). (c) MEMC (BB40044). (d) mUPGMA (BB40044).

The dense structure near the root can sometimes be beneficial for the BB4 category. The BB4 category requires introducing large external gaps in MSA results. The large external gap can be understood as a few point mutations that can significantly alter protein synthesis, such as early termination. Therefore, early bifurcations in trees may be reasonable for this category. For example, the MEMC algorithm produces the dense-root structure for the 44th test case in BB4 (BB40044) as shown in Fig. 4(c). The MSA process reflected this feature and provided a high amount of matches to the core blocks in the reference. In contrast, the mUPGMA guide trees did not capture these early bifurcations well (Fig. 4(d)). As a result, mUPGMA achieved a TC score of 0 in this example.

For simplicity, we collect the scores of the MEMC guide trees for `maxiterate 0, 1, 2`, and `1000` in Fig. 5 (see Appendix C for detailed results). As shown in Fig. 5, the MSA results using the MEMC trees were improved with the increasing number of iterations. Notably, the TC scores for BB3 were substantially improved from `maxiterate 0 to 1`. Even though the MEMC trees showed significantly lower TC scores for BB3 than the traditional trees, this can be overcome by only a single refinement step. This indicates that the MEMC guide tree does not locate the MSA tool at an incorrect initial point. Furthermore, these lower initial TC scores for BB3 may not affect MSA results in realistic situations where at least several iterations are necessary.

**FIG. 5.**
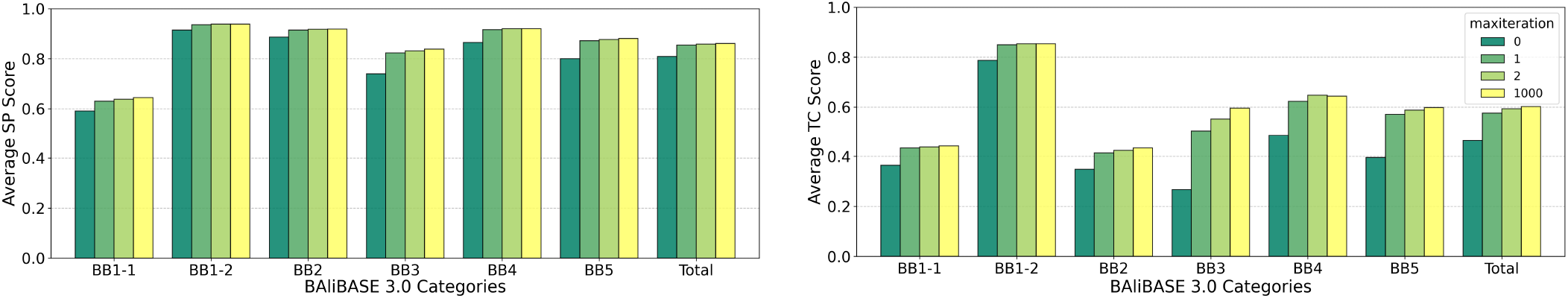
The SP and TC scores of MEMC guide trees are improved by iterative refinement. The most significant gains occurred from the first step.

## IV. DISCUSSION

The aim of the present work is to develop a quantum guide tree construction algorithm that can show better scalability than traditional methods. The MEMC algorithm theoretically requires *O*(*N*) computational time for a given distance matrix. This represents a quadratic improvement to the previous algorithms, which require *O*(*N*^2^).

The benchmark results in Section III were generated under the strict setting: only a single sampling was performed on the D-Wave hybrid solver for each test case. This condition was chosen to simulate a more realistic performance and prevent the overestimation by a large number of samplings. Even with this limitation, our method performed comparably to the classical methods in most cases.

The BB3 category was the only exception when no iteration was performed on MAFFT. The average TC score was significantly lower for this category, which may be due to the architecture of current quantum annealers. While the largest available D-Wave QPU has about 5,000 physical qubits, the qubits are not connected to all other qubits. This results in a significant overhead in the number of physical qubits required to solve a large TSP. The BB3 is the largest category on average; for instance, its largest TSP requires at least 142^2^ ≈ 20000 qubits. As this exceeds the capability of the current quantum annealers, the entire problem cannot be directly mapped onto qubits. Therefore, the hybrid solver must divide the problem into smaller parts by employing a classical heuristic method. Customizing a more efficient binary-to-qubit mapping can improve accuracy, but it is not an option in the hybrid solver. It is expected that these hardware limitations are temporary and will be overcome by ongoing advances in quantum hardware and algorithms. Additionally, a single refinement was enough to make the results of our method consistently comparable to the classical methods across all test categories.

We cannot exclude the possibility that the branch length estimation in our method may also have an impact on the results. Even if the MC has been largely accepted for phylogeny studies, it is based on the fossil findings of mammals. Bacteria, for example, have horizontal gene transfer mechanisms, which do not follow a constant molecular evolution. Therefore, a further study should focus on developing more accurate branch length estimation without significantly increasing computational complexity.

Surprisingly, the statistical analysis showed that the first iterative refinement was enough for the MEMC to achieve a nonsignificant score difference from the conventional methods. This finding implies that an MSA with a few iterations using a MEMC tree from a less accurate TSP solution can be enough to achieve a good MSA result. To enhance the runtime of our method, one can pre-terminate the quantum annealing process. Then, the less accurate TSP solution obtained can be directly used for constructing an MEMC tree.

However, the benchmark test should be carefully interpreted as an example, not as a rigorous demonstration of the performance. We selected the smallest benchmark test because of the size of the available quantum annealer. Further benchmark tests should be investigated, and different MSA tools should be tested.

It is well known that the accuracy of classical guide trees, including scalable algorithms (PartTree and mBed), quickly decreases as the problem size increases (see [85, Fig. 3]). Scalable guide tree algorithms such as PartTree are not recommended for small problems [86]. In contrast, the MEMC algorithm showed mostly comparable results with the classical methods in the small-scale benchmark test. While it was not tested here, the MEMC algorithm may surpass various classical guide tree algorithms in large-scale problems with the technical advances in quantum hardware.

## V. CONCLUSION

This work introduces a scalable guide tree construction algorithm that shows comparable performance to the classical methods by using only a single sampling in quantum annealing. We made a new interpretation of the ME and MC, referred to it as the MEMC principle, which considers an optimal tree to represent a minimum and constant rate of changes in sequences. It reduces the time complexity of determining positive branch lengths from *O*(*N*^2^) in the best implementations of the traditional algorithms to *O*(*N*).

The following two aspects provide the scalability of the MEMC algorithm. First, the ME problem is mapped to a TSP problem, which can be efficiently solved using quantum annealing. The greedy approach to the ME problem in classical tree algorithms is therefore avoided. Note that the MEMC algorithm can be implemented with classical TSP solvers for small problems, but it requires quantum annealing for larger instances. Secondly, an approximate TSP solution can serve as a good starting point to construct a guide tree, which permits achieving the linear time complexity.

For the BAliBASE 3.0 benchmark test, we limit the number of sampling in quantum annealing for each TSP to a single shot. The purpose was to avoid overestimating the performance of the MEMC algorithm with extensive sampling. Even with this strict setting, the SP scores of the MEMC trees were not significantly different from the classical ones statistically. The significantly low TC score of the BB3 category with no MSA refinements implies that further work is needed to improve the accuracy of the branch length estimation based on the MC. However, the low score may also be caused by the performance of the quantum hardware and the hybrid solver.

The benchmark test does not fully conclude the performance analysis. The best performance of the MEMC can be tested by increasing the number of samplings for quantum annealing. The MEMC algorithm is not confined to protein MSA; therefore, other large-scale MSA benchmark tests are required for a more comprehensive understanding of the algorithm. These tests will be possible in the near future as the quantum annealing hardware has been dramatically improved.

An interesting perspective on our guide tree algorithm is its applicability to general clustering problems. For instance, chemical similarity research [87] employs HAC such as Ward [88] to discover structure-activity relationships of chemicals based on their chemical structures. The MEMC algorithm may be useful in clustering chemicals that share a common structure with various additional functional groups, such as corticosteroid derivatives.

## AVAILABILITY OF DATA AND MATERIALS

The Python source code for the MEMC algorithm is provided via a GitHub repository (https://github.com/Bevacizumab/memcguidetree) under the BSD-3-Clause license. The BAliBASE 3.0 test sets were accessed from (https://www.lbgi.fr/balibase/BalibaseDownload/BAliBASE_R1-5.tar.gz) on July 29th, 2024. MAFFT (ver. 7.526) is available from a Gitlab repository: https://gitlab.com/sysimm/mafft/-/tree/ee9799916df6a5d5103d46d54933f8eb6d28e244. The raw data files generated for this study have been deposited in the Zenodo repository at https://doi.org/10.5281/zenodo.17115360. The summary of these results is available in the Appendix.

## ACKNOWLEDGEMENTS

Our thanks to Gwonhak Lee, You Kyoung Chung, Seon Bin Song, Jiyun Lee, and Kyoung An Park for insightful feedback on our work. This work was partly supported by Basic Science Research Program through the National Research Foundation of Korea (NRF), funded by the Ministry of Education, Science and Technology (2020M3H3A1110365, NRF-RS-2023-NR068116, RS-2023-NR119931, RS-2025-03532992, RS-2025-07882969). This work was also partly supported by Institute for Information & communications Technology Promotion (IITP) grant funded by the Korea government (MSIP) (No. 2019-0-00003, Research and Development of Core technologies for Programming, Running, Implementing and Validating of Fault-Tolerant Quantum Computing System). The Ministry of Trade, Industry, and Energy (MOTIE), Korea, also partly supported this research under the Industrial Innovation Infrastructure Development Project (Project No. RS-2024-00466693). JH is supported by the Yonsei University Research Fund of 2025-22-0140.

## APPENDIX Appendix A: Parallelizing Distance Matrix Computation

A simple method for the parallel computation of distance matrices is suggested here. First, a given set of sequences is sorted in increasing order of length because distance calculations for several long sequences can be more expensive than those for many short sequences. Calculating the distance between two sequences of lengths *m* and *n* requires *O*(*m* + *n*) and ≈ *O*(*mn*) time complexities for approximate measures [19, 89] and exact measures [90], respectively. As each sequence length is a positive integer with at most *k* digits, we can sort them in *O*(*kN*) ≈ *O*(*N*) time complexity using radix sort.

Let *s*_1_, *s*_2_, …, *s*_*N*_ be the *N* input sequences, sorted by length such that *s*_1_ is the shortest. Suppose that the number of threads available is *T*, which is smaller than or equal to *N* − 1. We define the following two values *q* and *r*:

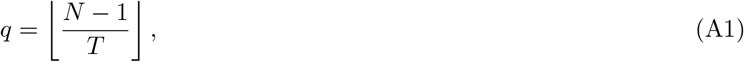

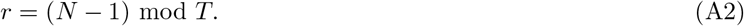

We also define *l*_*i*_ and *J*_*i*_, which are the labels of the first row assigned to the *i*-th thread and the set of distance computation jobs assigned to the *i*-th thread, respectively.

There are two cases: (1) *r* = 0, and (2) *r >* 0. When *r* = 0, each thread is equally assigned *q* number of rows:

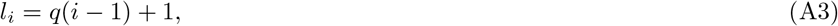

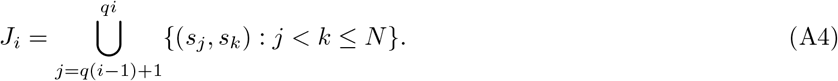

**FIG. A1.**
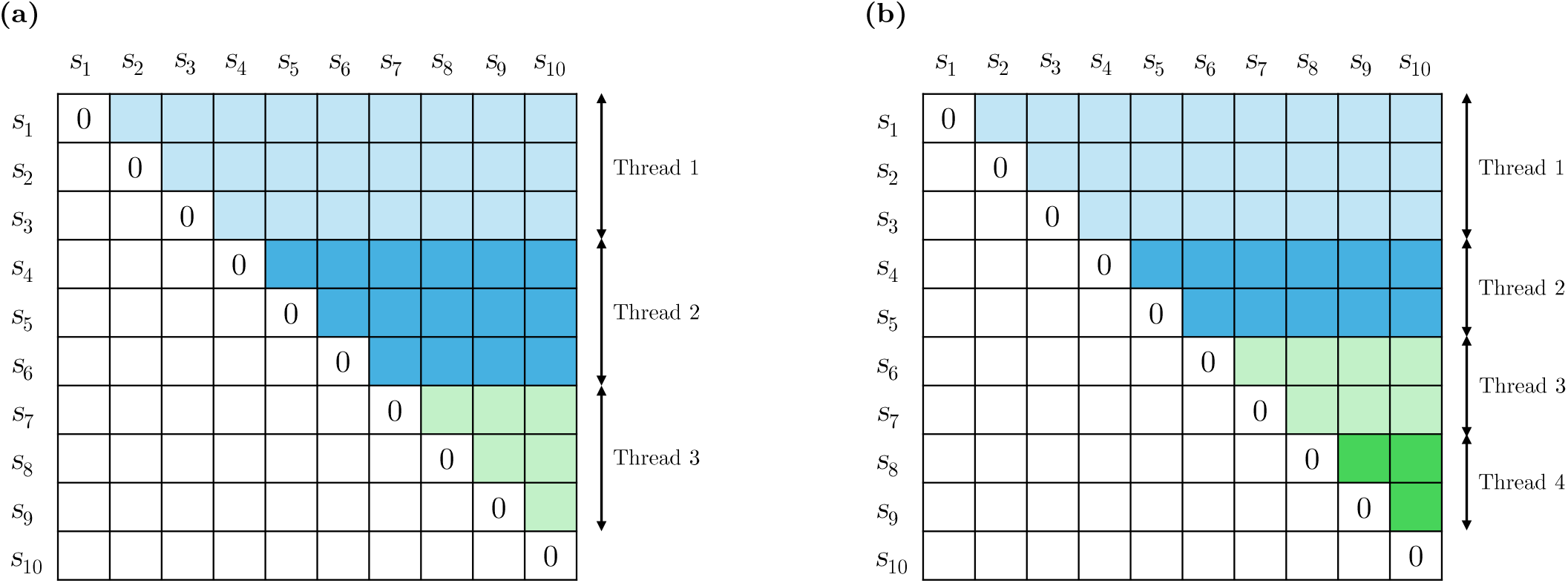
Examples of parallel computation of distance matrices. For a given set of 10 sequences, (a) uses three threads, and (b) uses four threads.

When *r >* 0, the first *r* threads are assigned *q* + 1 number of rows and the other threads have *q* number of rows:

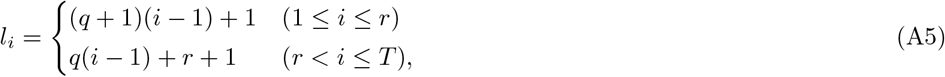

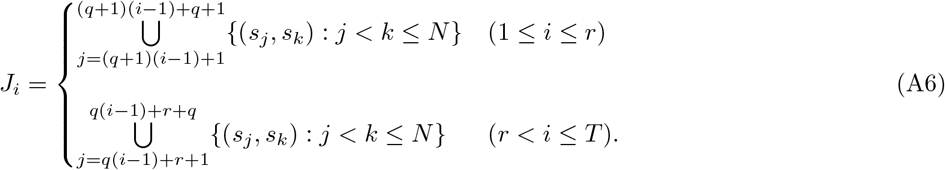

Fig. A1 shows the two cases for an example with 10 sequences: *r* = 0 and *r >* 0. The number of threads is 3 in Fig. A1(a), so *q* = 3 and *r* = 0. As *r* = 0, each thread is assigned an equal number of rows (*q* = 3) to compute. From Eq. A3, we obtain

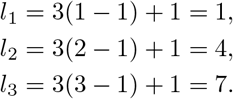

Thus, the second thread begins calculations from *s*_4_ (fourth row) to *s*_6_ (sixth row). Using Eq. A4, the set of jobs for the second thread is determined as follows:

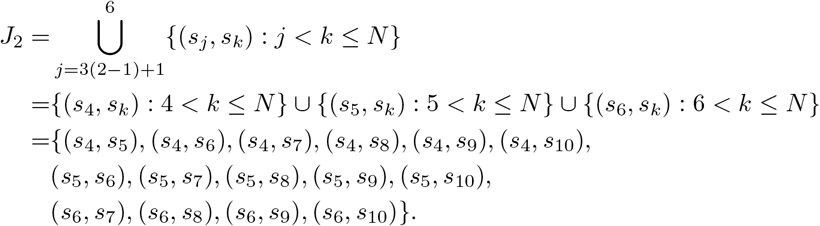

As shown in Fig. A1(a), Thread 1 processes the first three rows, which involves a lighter workload compared with Threads 2 and 3. Thus, the number of distance calculations assigned to Thread 1 is the largest, whereas Thread 3 has the smallest workload.

Fig. A1(b) shows when *q* = 2 and *r* = 1. If each thread is assigned two rows, one row will remain unassigned. Therefore, an additional row is assigned to Thread 1. By explicitly substituting *q* = 2 and *r* = 1, one can verify that Eq. A5 and Eq. A6 are correct.

This parallel computing scheme has the same time complexity as non-parallel computing, which scales quadratically with respect to the problem size. However, if *N* − 1 threads are used, this scheme can effectively reduce the runtime by a factor of *N* − 1.

## Appendix B: Data Preparation Details

### 1. BAliBASE 3.0: Protein MSA Benchmark

Only core blocks are considered for calculating SP and TC scores. Core blocks represent reliably aligned columns with no gaps in the MSA results [65]. They are determined based on whether three-dimensional structures are superposable and sequences are conserved at these sites.

BAliBASE provides MSA reference results indicating core blocks in XML format files. The core block data are located at the end of these files within the `<colsco-data>` tag. A value of 1 indicates a column that is determined to be a core block.

SP score is calculated by counting the number of pairwise matches to core blocks in a reference. TC score is computed by counting the number of columns that exactly match the core blocks in a reference. These scores range from 0 to 1, with higher scores indicating better alignments.

### 2. MAFFT L-INS-i

We obtain distance matrices from BAliBASE test cases using MAFFT L-INS-i:

~~~
mafft --localpair --maxiterate 0 --distout input_file
~~~

This command generates files with the `.hat2` extension that contains distance information in the input file directory. Note that the command does not specify a directory for the output file. By default, the code prints distances to three decimal places. To print more decimal places, the `WriteFloatHat2_pointer halfmtx` function in the file `io.c` has to be manually modified. This modification is limited to adjusting the number of decimal places printed and does not affect the performance of the program.

The MSA results are computed by the following command with default parameters:

~~~
mafft --localpair --maxiterate num --treein input_guide_tree_file input_file > output_file
~~~

where `num = 0, 1, 2` and `1000` for 0, 1, 2, and 1000 iterations, respectively.

### 3. D-Wave quantum annealer

A D-Wave quantum-classical hybrid solver (`hybrid_binary_quadratic_model_version2p`) was used to perform quantum annealing in the MEMC algorithm. The solver automatically embeds binary variables in the TSP to physical qubits; thus, we did not customize or optimize the embedding. TSPs up to 200 cities can be solved by the hybrid solver. As the largest number of sequences in the BAliBASE test cases is 142, the hybrid solver can solve all the TSPs in these benchmark tests.

## Appendix C: Supplementary Data

**TABLE C1.**
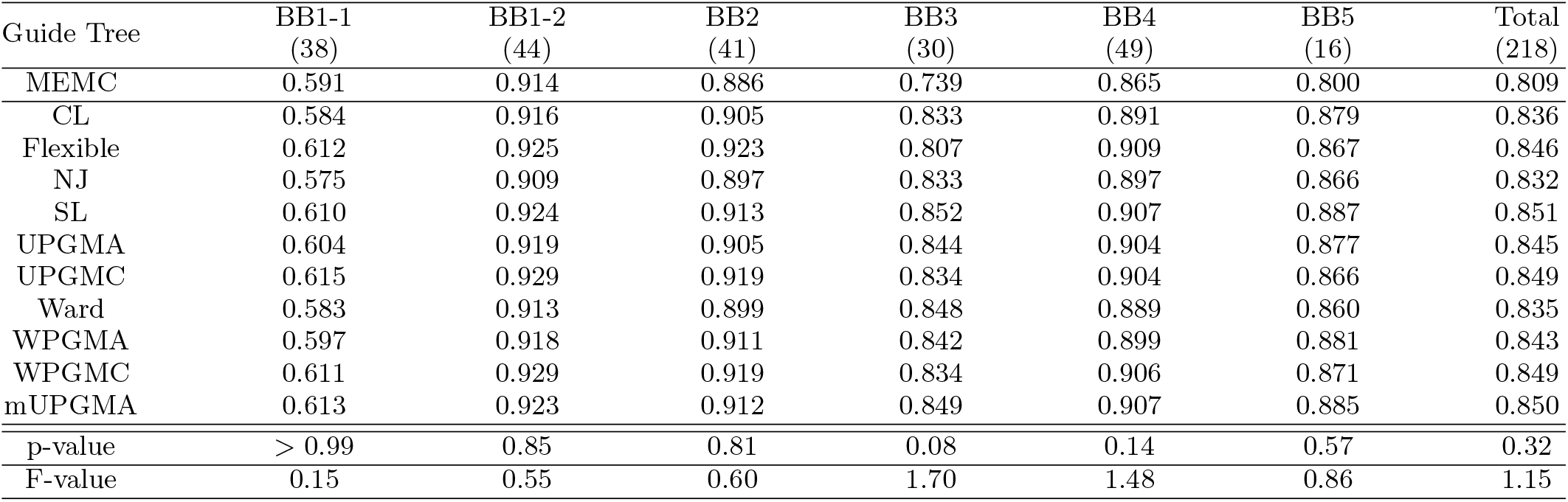
Average SP scores of guide tree algorithms for the BAliBASE 3.0 benchmark (`maxiterate 0`). The last column presents the average SP scores over all 218 benchmark test cases. The second row shows the benchmark test results when MEMC guide trees were used. The benchmark results using the classical guide tree algorithms are shown in alphabetical order. The last two rows give the p-values and F-values from one-way ANOVA tests of all guide trees for each column.

| Guide Tree | BB1-1<br>(38) | BB1-2<br>(44) | BB2<br>(41) | BB3<br>(30) | BB4<br>(49) | BB5<br>(16) | Total<br>(218) |
| --- | --- | --- | --- | --- | --- | --- | --- |
| MEMC | 0.591 | 0.914 | 0.886 | 0.739 | 0.865 | 0.800 | 0.809 |
| CL | 0.584 | 0.916 | 0.905 | 0.833 | 0.891 | 0.879 | 0.836 |
| Flexible | 0.612 | 0.925 | 0.923 | 0.807 | 0.909 | 0.867 | 0.846 |
| NJ | 0.575 | 0.909 | 0.897 | 0.833 | 0.897 | 0.866 | 0.832 |
| SL | 0.610 | 0.924 | 0.913 | 0.852 | 0.907 | 0.887 | 0.851 |
| UPGMA | 0.604 | 0.919 | 0.905 | 0.844 | 0.904 | 0.877 | 0.845 |
| UPGMC | 0.615 | 0.929 | 0.919 | 0.834 | 0.904 | 0.866 | 0.849 |
| Ward | 0.583 | 0.913 | 0.899 | 0.848 | 0.889 | 0.860 | 0.835 |
| WPGMA | 0.597 | 0.918 | 0.911 | 0.842 | 0.899 | 0.881 | 0.843 |
| WPGMC | 0.611 | 0.929 | 0.919 | 0.834 | 0.906 | 0.871 | 0.849 |
| mUPGMA | 0.613 | 0.923 | 0.912 | 0.849 | 0.907 | 0.885 | 0.850 |
| p-value | > 0.99 | 0.85 | 0.81 | 0.08 | 0.14 | 0.57 | 0.32 |
| F-value | 0.15 | 0.55 | 0.60 | 1.70 | 1.48 | 0.86 | 1.15 |

**TABLE C2.** Average TC scores of guide tree algorithms for the BAliBASE 3.0 benchmark (maxiterate 0).

| Guide Tree | BB1-1<br>(38) | BB1-2<br>(44) | BB2<br>(41) | BB3<br>(30) | BB4<br>(49) | BB5<br>(16) | Total<br>(218) |
| --- | --- | --- | --- | --- | --- | --- | --- |
| MEMC | 0.364 | 0.787 | 0.348 | 0.267 | 0.486 | 0.395 | 0.463 |
| CL | 0.354 | 0.796 | 0.388 | 0.586 | 0.543 | 0.548 | 0.538 |
| Flexible | 0.401 | 0.824 | 0.413 | 0.562 | 0.595 | 0.567 | 0.566 |
| NJ | 0.337 | 0.777 | 0.315 | 0.584 | 0.526 | 0.525 | 0.512 |
| SL | 0.372 | 0.819 | 0.432 | 0.605 | 0.569 | 0.581 | 0.565 |
| UPGMA | 0.375 | 0.803 | 0.414 | 0.598 | 0.564 | 0.554 | 0.555 |
| UPGMC | 0.401 | 0.833 | 0.426 | 0.610 | 0.569 | 0.563 | 0.571 |
| Ward | 0.326 | 0.801 | 0.350 | 0.597 | 0.528 | 0.524 | 0.523 |
| WPGMA | 0.368 | 0.801 | 0.413 | 0.591 | 0.556 | 0.558 | 0.551 |
| WPGMC | 0.400 | 0.833 | 0.421 | 0.587 | 0.585 | 0.567 | 0.571 |
| mUPGMA | 0.373 | 0.815 | 0.452 | 0.606 | 0.582 | 0.584 | 0.572 |
| p-value | 0.98 | 0.54 | 0.54 | < 0.001 | 0.91 | 0.84 | 0.003 |
| F-value | 0.31 | 0.90 | 0.89 | 5.02 | 0.47 | 0.57 | 2.69 |

**TABLE C3.** Average SP scores of guide tree algorithms for the BAliBASE 3.0 benchmark (maxiterate 1).

| Guide Tree | BB1-1<br>(38) | BB1-2<br>(44) | BB2<br>(41) | BB3<br>(30) | BB4<br>(49) | BB5<br>(16) | Total<br>(218) |
| --- | --- | --- | --- | --- | --- | --- | --- |
| MEMC | 0.630 | 0.936 | 0.914 | 0.823 | 0.916 | 0.872 | 0.854 |
| CL | 0.639 | 0.934 | 0.925 | 0.856 | 0.912 | 0.895 | 0.862 |
| Flexible | 0.641 | 0.937 | 0.923 | 0.818 | 0.919 | 0.886 | 0.859 |
| NJ | 0.642 | 0.935 | 0.923 | 0.866 | 0.925 | 0.899 | 0.867 |
| SL | 0.645 | 0.935 | 0.924 | 0.858 | 0.918 | 0.902 | 0.866 |
| UPGMA | 0.647 | 0.936 | 0.916 | 0.865 | 0.923 | 0.894 | 0.866 |
| UPGMC | 0.641 | 0.936 | 0.923 | 0.846 | 0.914 | 0.878 | 0.861 |
| Ward | 0.633 | 0.930 | 0.922 | 0.853 | 0.915 | 0.891 | 0.860 |
| WPGMA | 0.647 | 0.936 | 0.923 | 0.865 | 0.916 | 0.896 | 0.866 |
| WPGMC | 0.638 | 0.935 | 0.924 | 0.842 | 0.915 | 0.884 | 0.860 |
| mUPGMA | 0.640 | 0.935 | 0.924 | 0.860 | 0.918 | 0.901 | 0.865 |
| p-value | > 0.99 | > 0.99 | > 0.99 | 0.91 | > 0.99 | > 0.99 | > 0.99 |
| F-value | 0.018 | 0.051 | 0.076 | 0.47 | 0.17 | 0.18 | 0.14 |

**TABLE C4.** Average TC scores of guide tree algorithms for the BAliBASE 3.0 benchmark (maxiterate 1).

| Guide Tree | BB1-1<br>(38) | BB1-2<br>(44) | BB2<br>(41) | BB3<br>(30) | BB4<br>(49) | BB5<br>(16) | Total<br>(218) |
| --- | --- | --- | --- | --- | --- | --- | --- |
| MEMC | 0.434 | 0.850 | 0.414 | 0.504 | 0.623 | 0.571 | 0.576 |
| CL | 0.433 | 0.845 | 0.466 | 0.644 | 0.592 | 0.599 | 0.599 |
| Flexible | 0.450 | 0.852 | 0.450 | 0.592 | 0.639 | 0.605 | 0.605 |
| NJ | 0.438 | 0.850 | 0.454 | 0.677 | 0.634 | 0.605 | 0.614 |
| SL | 0.438 | 0.845 | 0.470 | 0.651 | 0.626 | 0.622 | 0.611 |
| UPGMA | 0.443 | 0.849 | 0.463 | 0.668 | 0.622 | 0.599 | 0.611 |
| UPGMC | 0.443 | 0.849 | 0.448 | 0.619 | 0.605 | 0.603 | 0.598 |
| Ward | 0.435 | 0.840 | 0.471 | 0.644 | 0.598 | 0.599 | 0.601 |
| WPGMA | 0.444 | 0.849 | 0.464 | 0.657 | 0.621 | 0.597 | 0.610 |
| WPGMC | 0.450 | 0.851 | 0.448 | 0.618 | 0.618 | 0.608 | 0.603 |
| mUPGMA | 0.440 | 0.845 | 0.471 | 0.655 | 0.626 | 0.619 | 0.612 |
| p-value | > 0.99 | > 0.99 | > 0.99 | 0.30 | > 0.99 | > 0.99 | 0.99 |
| F-value | 0.017 | 0.045 | 0.14 | 1.18 | 0.098 | 0.033 | 0.28 |

**TABLE C5.** Average SP scores of guide tree algorithms for the BAliBASE 3.0 benchmark (maxiterate 2).

| Guide Tree | BB1-1<br>(38) | BB1-2<br>(44) | BB2<br>(41) | BB3<br>(30) | BB4<br>(49) | BB5<br>(16) | Total<br>(218) |
| --- | --- | --- | --- | --- | --- | --- | --- |
| MEMC | 0.638 | 0.938 | 0.917 | 0.831 | 0.920 | 0.877 | 0.858 |
| CL | 0.642 | 0.936 | 0.926 | 0.857 | 0.913 | 0.898 | 0.864 |
| Flexible | 0.643 | 0.938 | 0.924 | 0.823 | 0.920 | 0.889 | 0.860 |
| NJ | 0.655 | 0.938 | 0.925 | 0.874 | 0.928 | 0.904 | 0.873 |
| SL | 0.648 | 0.936 | 0.926 | 0.861 | 0.920 | 0.903 | 0.867 |
| UPGMA | 0.644 | 0.939 | 0.917 | 0.862 | 0.926 | 0.897 | 0.867 |
| UPGMC | 0.642 | 0.936 | 0.925 | 0.848 | 0.915 | 0.880 | 0.862 |
| Ward | 0.645 | 0.934 | 0.924 | 0.854 | 0.924 | 0.894 | 0.866 |
| WPGMA | 0.645 | 0.938 | 0.924 | 0.864 | 0.918 | 0.897 | 0.867 |
| WPGMC | 0.638 | 0.936 | 0.925 | 0.842 | 0.915 | 0.886 | 0.861 |
| mUPGMA | 0.644 | 0.937 | 0.926 | 0.860 | 0.919 | 0.902 | 0.866 |
| p-value | >0.99 | >0.99 | >0.99 | 0.94 | 0.98 | >0.99 | >0.99 |
| F-value | 0.014 | 0.033 | 0.068 | 0.40 | 0.29 | 0.16 | 0.14 |

**TABLE C6.** Average TC scores of guide tree algorithms for the BAliBASE 3.0 benchmark (maxiterate 2).

| Guide Tree | BB1-1<br>(38) | BB1-2<br>(44) | BB2<br>(41) | BB3<br>(30) | BB4<br>(49) | BB5<br>(16) | Total<br>(218) |
| --- | --- | --- | --- | --- | --- | --- | --- |
| MEMC | 0.438 | 0.854 | 0.424 | 0.552 | 0.648 | 0.588 | 0.593 |
| CL | 0.441 | 0.851 | 0.475 | 0.654 | 0.596 | 0.609 | 0.607 |
| Flexible | 0.453 | 0.854 | 0.456 | 0.601 | 0.641 | 0.609 | 0.608 |
| NJ | 0.452 | 0.856 | 0.461 | 0.686 | 0.649 | 0.627 | 0.625 |
| SL | 0.440 | 0.850 | 0.478 | 0.657 | 0.630 | 0.628 | 0.616 |
| UPGMA | 0.447 | 0.858 | 0.476 | 0.668 | 0.639 | 0.613 | 0.621 |
| UPGMC | 0.443 | 0.848 | 0.456 | 0.629 | 0.606 | 0.608 | 0.602 |
| Ward | 0.449 | 0.848 | 0.486 | 0.654 | 0.621 | 0.616 | 0.615 |
| WPGMA | 0.453 | 0.855 | 0.463 | 0.663 | 0.626 | 0.605 | 0.615 |
| WPGMC | 0.453 | 0.851 | 0.455 | 0.617 | 0.619 | 0.610 | 0.605 |
| mUPGMA | 0.444 | 0.849 | 0.485 | 0.656 | 0.630 | 0.621 | 0.617 |
| p-value | >0.99 | >0.99 | >0.99 | 0.70 | >0.99 | >0.99 | >0.99 |
| F-value | 0.016 | 0.046 | 0.16 | 0.73 | 0.13 | 0.023 | 0.21 |

**TABLE C7.** Average SP scores of guide tree algorithms for the BAliBASE 3.0 benchmark (maxiterate 1000).

| Guide Tree | BB1-1<br>(38) | BB1-2<br>(44) | BB2<br>(41) | BB3<br>(30) | BB4<br>(49) | BB5<br>(16) | Total<br>(218) |
| --- | --- | --- | --- | --- | --- | --- | --- |
| MEMC | 0.644 | 0.938 | 0.918 | 0.838 | 0.920 | 0.881 | 0.861 |
| CL | 0.649 | 0.936 | 0.926 | 0.858 | 0.913 | 0.900 | 0.866 |
| Flexible | 0.646 | 0.938 | 0.924 | 0.826 | 0.922 | 0.890 | 0.862 |
| NJ | 0.664 | 0.938 | 0.927 | 0.881 | 0.930 | 0.906 | 0.876 |
| SL | 0.651 | 0.936 | 0.927 | 0.861 | 0.920 | 0.903 | 0.868 |
| UPGMA | 0.649 | 0.938 | 0.918 | 0.863 | 0.927 | 0.900 | 0.868 |
| UPGMC | 0.643 | 0.937 | 0.926 | 0.850 | 0.915 | 0.883 | 0.863 |
| Ward | 0.648 | 0.934 | 0.925 | 0.856 | 0.926 | 0.896 | 0.867 |
| WPGMA | 0.650 | 0.939 | 0.926 | 0.866 | 0.919 | 0.900 | 0.869 |
| WPGMC | 0.640 | 0.936 | 0.926 | 0.842 | 0.916 | 0.887 | 0.861 |
| mUPGMA | 0.648 | 0.937 | 0.927 | 0.861 | 0.920 | 0.903 | 0.868 |
| p-value | >0.99 | >0.99 | >0.99 | 0.95 | 0.97 | >0.99 | >0.99 |
| F-value | 0.027 | 0.030 | 0.065 | 0.39 | 0.33 | 0.15 | 0.17 |

**TABLE C8.** Average TC scores of guide tree algorithms for the BAliBASE 3.0 benchmark (maxiterate 1000).

| Guide Tree | BB1-1<br>(38) | BB1-2<br>(44) | BB2<br>(41) | BB3<br>(30) | BB4<br>(49) | BB5<br>(16) | Total<br>(218) |
| --- | --- | --- | --- | --- | --- | --- | --- |
| MEMC | 0.442 | 0.854 | 0.434 | 0.596 | 0.644 | 0.598 | 0.602 |
| CL | 0.448 | 0.854 | 0.470 | 0.654 | 0.598 | 0.616 | 0.608 |
| Flexible | 0.447 | 0.854 | 0.457 | 0.588 | 0.641 | 0.600 | 0.605 |
| NJ | 0.477 | 0.856 | 0.460 | 0.705 | 0.652 | 0.633 | 0.633 |
| SL | 0.451 | 0.849 | 0.490 | 0.663 | 0.630 | 0.632 | 0.621 |
| UPGMA | 0.450 | 0.857 | 0.479 | 0.666 | 0.641 | 0.622 | 0.623 |
| UPGMC | 0.450 | 0.848 | 0.458 | 0.627 | 0.609 | 0.607 | 0.603 |
| Ward | 0.452 | 0.850 | 0.497 | 0.657 | 0.627 | 0.616 | 0.620 |
| WPGMA | 0.452 | 0.858 | 0.474 | 0.663 | 0.627 | 0.625 | 0.619 |
| WPGMC | 0.460 | 0.850 | 0.463 | 0.617 | 0.622 | 0.613 | 0.608 |
| mUPGMA | 0.451 | 0.851 | 0.490 | 0.656 | 0.630 | 0.623 | 0.620 |
| p-value | >0.99 | >0.99 | >0.99 | 0.80 | >0.99 | >0.99 | >0.99 |
| F-value | 0.042 | 0.040 | 0.17 | 0.61 | 0.12 | 0.026 | 0.24 |

## References

[1] F. Sievers, A. Wilm, D. Dineen, T. J. Gibson, K. Karplus, W. Li, R. Lopez, H. McWilliam, M. Remmert, J. Söding, et al., Fast, scalable generation of high-quality protein multiple sequence alignments using clustal omega, Mol. Syst. Biol. 7, 539 (2011).

[2] R. C. Edgar and S. Batzoglou, Multiple sequence alignment, Curr. Opin. Struct. Biol. 16, 368 (2006).

[3] D. N. Gilbert et al., The Sanford guide to antimicrobial therapy (Antimicrobial Therapy, Inc., Sperryville, 2023).

[4] K. Takeda, S. Sakakibara, K. Yamashita, D. Motooka, S. Nakamura, M. A. El Hussien, J. Katayama, Y. Maeda, M. Nakata, S. Hamada, et al., Allergic conversion of protective mucosal immunity against nasal bacteria in patients with chronic rhinosinusitis with nasal polyposis, J. Allergy Clin. Immunol. 143, 1163 (2019).

[5] C. L. Pierri, G. Parisi, and V. Porcelli, Computational approaches for protein function prediction: a combined strategy from multiple sequence alignment to molecular docking-based virtual screening, BBA-Proteins Proteomics 1804, 1695 (2010).

[6] D. P. Ryan and J. M. Matthews, Protein–protein interactions in human disease, Curr. Opin. Struct. Biol. 15, 441 (2005).

[7] D. Collias and C. L. Beisel, Crispr technologies and the search for the pam-free nuclease, Nat. Commun. 12, 555 (2021).

[8] A. W. Senior, R. Evans, J. Jumper, J. Kirkpatrick, L. Sifre, T. Green, C. Qin, A. Žídek, A. W. R. Nelson, A. Bridgland, et al., Improved protein structure prediction using potentials from deep learning, Nature 577, 706 (2020).

[9] J. Jumper, R. Evans, A. Pritzel, T. Green, M. Figurnov, O. Ronneberger, K. Tunyasuvunakool, R. Bates, A. Žídek, A. Potapenko, et al., Highly accurate protein structure prediction with alphafold, Nature 596, 583 (2021).

[10] M. Ekeberg, C. Lövkvist, Y. Lan, M. Weigt, and E. Aurell, Improved contact prediction in proteins: using pseudolikelihoods to infer potts models, Phys. Rev. E 87, 012707 (2013).

[11] W. Just, Computational complexity of multiple sequence alignment with sp-score, J. Comput. Biol. 8, 615 (2001).

[12] L. Wang and T. Jiang, On the complexity of multiple sequence alignment, J. Comput. Biol. 1, 337 (1994).

[13] A. Löytynoja, A. J. Vilella, and N. Goldman, Accurate extension of multiple sequence alignments using a phylogeny-aware graph algorithm, Bioinformatics 28, 1684 (2012).

[14] A. Löytynoja and N. Goldman, An algorithm for progressive multiple alignment of sequences with insertions, Proc. Natl. Acad. Sci. U.S.A. 102, 10557 (2005).

[15] A. Loytynoja and N. Goldman, Phylogeny-aware gap placement prevents errors in sequence alignment and evolutionary analysis, Science 320, 1632 (2008).

[16] M. A. Larkin, G. Blackshields, N. P. Brown, R. Chenna, P. A. McGettigan, H. McWilliam, F. Valentin, I. M. Wallace, A. Wilm, R. Lopez, et al., Clustal w and clustal x version 2.0, Bioinformatics 23, 2947 (2007).

[17] C. Grasso and C. Lee, Combining partial order alignment and progressive multiple sequence alignment increases alignment speed and scalability to very large alignment problems, Bioinformatics 20, 1546 (2004).

[18] B. Morgenstern, K. Frech, A. Dress, and T. Werner, Dialign: finding local similarities by multiple sequence alignment, Bioinformatics 14, 290 (1998).

[19] T. Lassmann and E. L. Sonnhammer, Kalign–an accurate and fast multiple sequence alignment algorithm, BMC Bioinformatics 6, 298 (2005).

[20] K. Katoh and D. M. Standley, Mafft multiple sequence alignment software version 7: improvements in performance and usability, Mol. Biol. Evol. 30, 772 (2013).

[21] Y. Liu, B. Schmidt, and D. L. Maskell, Msaprobs: multiple sequence alignment based on pair hidden markov models and partition function posterior probabilities, Bioinformatics 26, 1958 (2010).

[22] R. C. Edgar, Muscle: multiple sequence alignment with high accuracy and high throughput, Nucleic Acids Res. 32, 1792 (2004).

[23] R. C. Edgar, Muscle: a multiple sequence alignment method with reduced time and space complexity, BMC Bioinformatics 5, 113 (2004).

[24] U. Roshan and D. R. Livesay, Probalign: multiple sequence alignment using partition function posterior probabilities, Bioinformatics 22, 2715 (2006).

[25] C. B. Do, M. S. P. Mahabhashyam, M. Brudno, and S. Batzoglou, Probcons: Probabilistic consistency-based multiple sequence alignment, Genome Res. 15, 330 (2005).

[26] C. Notredame, D. G. Higgins, and J. Heringa, T-coffee: A novel method for fast and accurate multiple sequence alignment, J. Mol. Biol. 302, 205 (2000).

[27] S. B. Needleman and C. D. Wunsch, A general method applicable to the search for similarities in the amino acid sequence of two proteins, J. Mol. Biol. 48, 443 (1970).

[28] T. F. Smith, M. S. Waterman, et al., Identification of common molecular subsequences, J. Mol. Biol. 147, 195 (1981).

[29] D.-F. Feng and R. F. Doolittle, Progressive sequence alignment as a prerequisitetto correct phylogenetic trees, J. Mol. Evol. 25, 351 (1987).

[30] M. Chatzou, C. Magis, J.-M. Chang, C. Kemena, G. Bussotti, I. Erb, and C. Notredame, Multiple sequence alignment modeling: methods and applications, Brief. Bioinform. 17, 1009 (2016).

[31] A. Stamatakis, Raxml-vi-hpc: maximum likelihood-based phylogenetic analyses with thousands of taxa and mixed models, Bioinformatics 22, 2688 (2006).

[32] Z. Yang, Phylogenetic analysis using parsimony and likelihood methods, J. Mol. Evol. 42, 294 (1996).

[33] J. P. Huelsenbeck, B. Larget, R. E. Miller, and F. Ronquist, Potential applications and pitfalls of bayesian inference of phylogeny, Syst. Biol. 51, 673 (2002).

[34] Q. Zhan, Y. Ye, T.-W. Lam, S.-M. Yiu, Y. Wang, and H.-F. Ting, Improving multiple sequence alignment by using better guide trees, BMC Bioinformatics 16, S4 (2015).

[35] R. Agarwala, V. Bafna, M. Farach, M. Paterson, and M. Thorup, On the approximability of numerical taxonomy (fitting distances by tree metrics), SIAM J. Comput. 28, 1073 (1998).

[36] D. Catanzaro, M. Labbé, R. Pesenti, and J.-J. Salazar-González, Mathematical models to reconstruct phylogenetic trees under the minimum evolution criterion, Networks 53, 126 (2009).

[37] W. H. E. Day, Computational complexity of inferring phylogenies from dissimilarity matrices, Bull. Math. Biol. 49, 461 (1987).

[38] D. Catanzaro, The minimum evolution problem: Overview and classification, Networks 53, 112 (2009).

[39] Y. Loewenstein, E. Portugaly, M. Fromer, and M. Linial, Efficient algorithms for accurate hierarchical clustering of huge datasets: tackling the entire protein space, Bioinformatics 24, i41 (2008).

[40] A. Krause, J. Stoye, and M. Vingron, Large scale hierarchical clustering of protein sequences, BMC Bioinformatics 6, 15 (2005).

[41] N. Saitou and M. Nei, The neighbor-joining method: a new method for reconstructing phylogenetic trees, Mol. Biol. Evol. 4, 406 (1987).

[42] R. R. Sokal and C. D. Michener, A statistical method for evaluating systematic relationships, Univ. Kansas Sci. Bull. 38, 1409 (1958).

[43] K. Howe, A. Bateman, and R. Durbin, Quicktree: building huge neighbour-joining trees of protein sequences, Bioinformatics 18, 1546 (2002).

[44] J. Evans, L. Sheneman, and J. Foster, Relaxed neighbor joining: a fast distance-based phylogenetic tree construction method, J. Mol. Evol. 62, 785 (2006).

[45] M. Simonsen, T. Mailund, and C. N. Pedersen, Rapid neighbour-joining, in Proceedings of the International Workshop on Algorithms in Bioinformatics (Springer, 2008) pp. 113–122.

[46] I. Elias and J. Lagergren, Fast neighbor joining, Theor. Comput. Sci. 410, 1993 (2009).

[47] K. Katoh and H. Toh, Parttree: an algorithm to build an approximate tree from a large number of unaligned sequences, Bioinformatics 23, 372 (2007).

[48] G. Blackshields, F. Sievers, W. Shi, A. Wilm, and D. G. Higgins, Sequence embedding for fast construction of guide trees for multiple sequence alignment, Algorithms. Mol. Biol. 5, 21 (2010).

[49] K. Nalecz-Charkiewicz and R. M. Nowak, Algorithm for dna sequence assembly by quantum annealing, BMC Bioinformatics 23, 122 (2022).

[50] P.-N. Nguyen, Biomarker discovery with quantum neural networks: a case-study in ctla4-activation pathways, BMC Bioinformatics 25, 149 (2024).

[51] S. Y. Madsen, F. K. Marqversen, S. E. Rasmussen, and N. T. Zinner, Multi-sequence alignment using the quantum approximate optimization algorithm, arXiv preprint (2023), arXiv:2308.12103.

[52] O. B. Lindvall, Quantum Methods for Sequence Alignment and Metagenomics, Ph.d. thesis, KTH Royal Institute of Technology (2019).

[53] M. Lee, Modified multiple sequence alignment algorithm on quantum annealers (maq), arXiv preprint (2024), arXiv:2403.17979.

[54] Y. K. Chung, D. Lee, J. Lee, J. Kim, D. K. Park, and J. Huh, Quantum-classical hybrid approach for codon optimization and its practical applications, bioRxiv preprint 10.1101/2024.06.08.598046 (2024).

[55] A. B. Finnila, M. A. Gomez, C. Sebenik, C. Stenson, and J. D. Doll, Quantum annealing: A new method for minimizing multidimensional functions, Chem. Phys. Lett. 219, 343 (1994).

[56] E. Farhi, J. Goldstone, and S. Gutmann, A quantum approximate optimization algorithm, arXiv preprint (2014), arXiv:1411.4028.

[57] A. Rajak, S. Suzuki, A. Dutta, and B. K. Chakrabarti, Quantum annealing: An overview, Phil. Trans. R. Soc. A. 381, 20210417 (2023).

[58] M. W. Johnson, M. H. S. Amin, S. Gildert, T. Lanting, F. Hamze, N. Dickson, R. Harris, A. J. Berkley, J. Johansson, P. Bunyk, et al., Quantum annealing with manufactured spins, Nature 473, 194 (2011).

[59] B. Bauer, S. Bravyi, M. Motta, and G. K.-L. Chan, Quantum algorithms for quantum chemistry and quantum materials science, Chemical reviews 120, 12685 (2020).

[60] C. Mc Keever and M. Lubasch, Towards adiabatic quantum computing using compressed quantum circuits, PRX quantum 5, 020362 (2024).

[61] M. Demirplak and S. A. Rice, Adiabatic population transfer with control fields, J. Phys. Chem. A 107, 9937 (2003).

[62] P. Díez-Valle, F. J. Gómez-Ruiz, D. Porras, and J. J. García-Ripoll, Universal resources for qaoa and quantum annealing, arXiv preprint (2025), arXiv:2506.03241.

[63] T. Albash and D. A. Lidar, Demonstration of a scaling advantage for a quantum annealer over simulated annealing, Phys. Rev. X 8, 031016 (2018).

[64] A. D. King, J. Raymond, T. Lanting, S. V. Isakov, M. Mohseni, G. Poulin-Lamarre, S. Ejtemaee, W. Bernoudy, I. Ozfidan, A. Y. Smirnov, et al., Scaling advantage over path-integral monte carlo in quantum simulation of geometrically frustrated magnets, Nat. Commun. 12, 1113 (2021).

[65] J. D. Thompson, P. Koehl, R. Ripp, and O. Poch, Balibase 3.0: latest developments of the multiple sequence alignment benchmark, Proteins 61, 127 (2005).

[66] A. Rzhetsky and M. Nei, A simple method for estimating and testing minimum-evolution trees, Mol. Biol. Evol. 9, 945 (1992).

[67] A. Rzhetsky and M. Nei, Theoretical foundation of the minimum-evolution method of phylogenetic inference, Mol. Biol. Evol. 10, 1073 (1993).

[68] O. Gascuel and M. Steel, Neighbor-joining revealed, Mol. Biol. Evol. 23, 1997 (2006).

[69] L. L. Cavalli-Sforza and A. W. Edwards, Phylogenetic analysis. models and estimation procedures, Am. J. Hum. Genet. 19, 233 (1967).

[70] A. Drummond and A. G. Rodrigo, Reconstructing genealogies of serial samples under the assumption of a molecular clock using serial-sample upgma, Mol. Biol. Evol. 17, 1807 (2000).

[71] S. Kumar, Molecular clocks: four decades of evolution, Nat. Rev. Genet. 6, 654 (2005).

[72] E. Zuckerkandl and L. Pauling, Molecular Disease, Evolution, and Genic Heterogeneity (Academic Press, Cambridge, 1962).

[73] E. Zuckerkandl and L. Pauling, Evolutionary divergence and convergence in proteins, in Evolving Genes and Proteins, edited by H. J. V. Vernon Bryson (Academic Press, Cambridge, 1965) pp. 97–166.

[74] C. Korostensky and G. H. Gonnet, Using traveling salesman problem algorithms for evolutionary tree construction, Bioinformatics 16, 619 (2000).

[75] C. Korostensky and G. Gonnet, Near optimal multiple sequence alignments using a traveling salesman problem approach, in Proceedings of the 6th International Symposium on String Processing and Information Retrieval (SPIRE’99) (IEEE, 1999) pp. 105–114.

[76] K. Giannakis, C. Papalitsas, G. Theocharopoulou, S. Fanarioti, and T. Andronikos, A quantum-inspired optimization heuristic for the multiple sequence alignment problem in bio-computing, in Proceedings of the 2019 10th International Conference on Information, Intelligence, Systems and Applications (IISA) (IEEE, 2019) pp. 1–8.

[77] A. Lucas, Ising formulations of many np problems, Front. Physics 2, 5 (2014).

[78] J. Choi and J. Kim, A tutorial on quantum approximate optimization algorithm (qaoa): Fundamentals and applications, in Proceedings of the 2019 international conference on information and communication technology convergence (ICTC) (IEEE, 2019) pp. 138–142.

[79] T. H. Cormen, C. E. Leiserson, R. L. Rivest, and C. Stein, Introduction to algorithms (MIT press, Cambridge, 2022).

[80] G. W. Milligan, Ultrametric hierarchical clustering algorithms, Psychometrika 44, 343 (1979).

[81] K. Katoh, J. Rozewicki, and K. D. Yamada, Mafft online service: multiple sequence alignment, interactive sequence choice and visualization, Brief. Bioinform. 20, 1160 (2019).

[82] G. N. Lance and W. T. Williams, A general theory of classificatory sorting strategies: 1. hierarchical systems, Comput. J. 9, 373 (1967).

[83] F. Sievers, G. M. Hughes, and D. G. Higgins, Systematic exploration of guide-tree topology effects for small protein alignments, BMC Bioinformatics 15, 338 (2014).

[84] J. D. Thompson, F. Plewniak, and O. Poch, A comprehensive comparison of multiple sequence alignment programs, Nucleic Acids Res. 27, 2682 (1999).

[85] K. Boyce, F. Sievers, and D. G. Higgins, Simple chained guide trees give high-quality protein multiple sequence alignments, Proc. Natl. Acad. Sci. U.S.A. 111, 10556 (2014).

[86] K. Katoh and H. Toh, Recent developments in the mafft multiple sequence alignment program, Brief. Bioinform. 9, 286 (2008).

[87] N. Nikolova and J. Jaworska, Approaches to measure chemical similarity–a review, QSAR Comb. Sci. 22, 1006 (2003).

[88] R. D. Brown and Y. C. Martin, Use of structure-activity data to compare structure-based clustering methods and descriptors for use in compound selection, J. Chem. Inf. Comput. Sci. 36, 572 (1996).

[89] S. Wu and U. Manber, Fast text searching: allowing errors, Commun. ACM 35, 83 (1992).

[90] S. R. Eddy, What is dynamic programming?, Nat. Biotechnol. 22, 909 (2004).

